# Immunoinformatics Prediction of Epitope Based Peptide Vaccine Against Schistosoma Mansoni Fructose Bisphosphate Aldolase Protein

**DOI:** 10.1101/755959

**Authors:** Mustafa Elhag, Ruaa Mohamed Alaagib, Esraa Musa Haroun, Nagla Mohamed Ahmed, Sahar Obi Abd Albagi, Mohammed A. Hassan

## Abstract

*Schistosoma Mansoni* represents an important tropical disease that can cause schistosomiasis mostly in Africa and Middle East with high mortality rates. Moreover, no vaccine against it exists. This study predicts an effective epitope-based vaccine against Fructose 1,6 Bisphosphate Aldolase (FBA) enzyme of Schistosoma Mansoni using immunoinformatics approaches. FBA is important for production of energy required for different schistosome activities and survival. The sequences were retrieved from NCBI and several prediction tests were conducted to analyze possible epitopes for B-cell, T-cell MHC class I and II. Tertiary structure of the most promising epitopes was obtained. Two epitopes showed high binding affinity for B-cells, while four epitopes showed high binding affinity for MHCI and MHCII. The results were promising to formulate a vaccine with more than 99.5% population coverage. We hope that these promising epitopes serves as a preventive measure for the disease in the future and recommend invivo and invitro studies.

## INTRODUCTION

*Schistosomiasis* or billharzias is one of the major neglected tropical diseases worldwide, ranking second only to malaria among parasitic diseases in terms of its socioeconomic and public health importance in tropical and subtropical areas affecting 200 million individuals in 74 countries[1]. The disease is caused by an infection with blood flukes of the genous Shistosoma [2]. Acute Schistosomiasis (also called Katayama fever or syndrome) occurs several weeks after the penetration of schistosome cercariae through the skin, and is related to the migration of the larvae (schistosomulae) within the body [3]. Considering geographical distributions, there are different Schistosoma species that can infect humans. S. mansoni and S. hematobium are mostly endemic to Africa and Middle East, which represents about 85% of the reported world cases [2]. S. mansoni is also found in South and Central America [4].

*S. Mansoni* inhabits the intestinal venules (in close contact with host humoral and cellular cytotoxic factors) and primarily affects the liver and gut [5, 6]. The infectious phase (cercariae) is developed particularly in fresh water areas, where the snails carrying the Schistosoma sporocytes exist. Cercariae penetrate the human skin and then migrate via blood circulation to the portal vein where the female parasite produces eggs. This leads to the appearance of acute and chronic symptoms such as fever, abdominal and anemia pain, exercise intolerance, and bloody diarrhea, [7] many eggs get trapped in organs such as the liver, where they induce a granulomatous inflammation and organ damage, the main cause of pathology in schistosomiasis [8]. Worldwide, 10% of the intestinal schistosomiasis, S. mansoni infection is associated with hepatic periportal fibrosis, hepatosplenomegaly, and esophageal varices as a result of long term exposure to the highly antigenic eggs [9]. Heavily infected individuals often develop severe morbidity with hepatosplenomegaly, sometimes with a fatal outcome. Morbidity is attributed to the strong humoral and T-cell-mediated host immune responses developed to a variety of parasite antigens and expressed as tissue inflammations [10].

This disease presents a substantial public health and economic burden and is considered a disease of poverty. An estimated 779 million people are at risk of infection, and approximately 252 million people are currently infected [11, 12]. The Global Health Estimates of 2015 attributed 3.51 million disability-adjusted life years (DALYs) and 10.1 million deaths in 2016 to *Schistosomiasis*, which is a mortality figure that has been challenged as a gross underestimate [13, 14]. Therefore, the WHO aims to eliminate *Schistosomiasis* as a public health problem globally by 2025 [15].

For the past 40 years, Praziquantel (PZQ) has been recommended by the World Health Organization for the treatment of all forms of *Schistosomiasis* [16]. On the other hand, the dependence on PZQ raises legitimate concerns about the appearance of drug resistance [17]. Although widespread resistance has not been convincingly demonstrated, field and experimental isolates displaying reduced sensitivity to PZQ have been described from several countries [18]. The discovery and development of novel effective drugs, and vaccine have been considered a research priority. Effective control of Schistosomiasis is unlikely in the absence of a vaccine [19]. Previously, we have identified the *S. Mansoni* glycolytic enzyme fructose-1, 6-bisphosphate aldolase (FBA) cDNA (1,431 base pair) clone (SMALDO), which has been recognized by protective antibodies taken from rabbits vaccinated with irradiated cercariae. [20] Sera of these rabbits were able to passively transfer high levels of resistance against *S. Mansoni* challenge infection to naïve mice when it was given around the time of the challenge. Carbohydrate metabolism in schistosomes points to a dominant anaerobic metabolism through glycolysis in vertebrate-dwelling stages of the parasite [21]. FBA plays a central role in glycolysis by catalyzing the reversible aldol cleavage of Fructose Bisphosphate into the two trioses, dihydroxyacetone-phosphate and glyceraldehyde-3-phosphate, [22] and it is important for production of energy required for different schistosome activities and survival. [23]. It contains the regions typical of eukaryotic mRNA. FBA which forms a Schiff’s base between the substrate and a lysyl residue, belongs to class I aldolases [24] whereas class II aldolases use a divalent ion as a cofactor. It has 61-70% sequence homology with other FBP aldolases from different species and higher vertebrates (type A aldolase) [20]. It is highly expressed in different life cycle stages, [20] and is a potentially suitable target for intervention. Fructose-1,6-bisphosphate aldolase have a significant role in pathogenesis by interacting with host’s antigene [25]. Knowing of epitope/antibody interaction is the key to construct potent vaccines and effective diagnostics [26]. Selected epitopes are capable of inducing B cell and T cell mediated immunity. This enzyme also induces strong humoral and cell-mediated immune responses and has protective and granuloma-modulating effects in animals vaccinated with the recombinant protein [27, 28]. Most of immunoinformatics researches are stressed on the design and study of algorithms for mapping potential B- and T-cell epitopes that speed up the time and lower the costs needed for laboratory analysis of pathogen products. Using such tools and information (reverse vaccinology) to analyse the sequence areas with potential binding sites can lead to the development of new vaccines [29].

The aim of this study is to predict peptide based vaccine from the highly conserved candidate protein FBA of *S. Mansoni* using computational approach. Many studies showed the immunological efficacy of peptide-based vaccines against infectious diseases. The development of peptide-based vaccines has significantly advanced with the identification of specific epitopes derived from infectious pathogens. Understanding of the molecular basis of antigen recognition and HLA binding motifs has resulted in the development of designed vaccines based on motifs predicted to bind to host class I or class II MHC [30]. No previous reports were found in *Schistosoma Mansoni* epitope based vaccine so this may be considered the first study to use insilico approach to design an epitope-based vaccine.

## MATERIALS AND METHODS

### Protein Sequence Retrieval

A total of four *Schistosoma mansoni* Fructose-bisphosphate aldolase protein sequences were obtained from NCBI (www.ncbi.nlm.nih.gov/) in FASTA format on 10/7/2019 (Table 1). The locations of continuous epitopes have been correlated with various parameters (such as hydrophobicity, flexibility, accessibility, turns, exposed surface, polarity, and antigenic propensity of polypeptide chain). The epitope predictions are based on propensity scales for each of the 363 amino acids.

**Table 1:** Countries and accession numbers of retrieved sequences from NCBI.

| Region | Country | Accession number | Protein |
| --- | --- | --- | --- |
| - | N.A | P53442.1 | Fructose- bisphosphate aldolase |
| - | UK | CCD79690.1 |  |
| - | UK | XP_018652294.1 |  |
| - | N.A | AAA57567.1 |  |

### Determination of conserved regions

All sequences of each protein were subjected to multiple sequence alignments using CLUSTALW tool of BIOEDIT sequence alignment editor (version 7.2.5.0) in order to identify conserved regions between sequences. Then epitopes prediction and analysis of each protein was done using different tools of immune epitope data base IEBD software (http://www.iedb.org) [31, 32].

### Sequenced-Based Method

The reference sequence (XP_018652294.1) of *Schistosoma Mansoni* FBA was subjected to different prediction tools at the Immune Epitope Database (IEDB) analysis resource (http://www.iedb.org/) to predict various B and T cell epitopes. Conserved epitopes would be considered as candidate epitopes for B and T cells [31].

### Prediction of B cell epitopes

The initial step when identifying antigenic epitopes parts in our target pathogens is the identification of linear peptide parts. A combination between (Parker and Levitt) method and hidden Markov model (HMM) were used to predict epitopes precisely [33]. Using IEDB software mainly Bepipred linear epitope prediction tool, B cell epitopes from conserved regions were identified from the protein with specific default threshold value for each protein [34].

Surface accessibility prediction: The calculate of surface probability method were using to increase sureness in provisional alignment for relating with other sequences was predicted by the garnier et al. Method chou and fasman method, Also this surface probability method accepts the absence of significant internal deletions or insertions. Emini surface accessibility prediction tool is based on Emini’s surface accessibility scale. Accessibility profile is predicted using the formula Sn = (i-1Π6δn + 4 + i) × (0.37)-6 where Sn is the surface probability, δn is the fractional surface probability value, and i vary from1 to 6. A hexapeptide sequence with Sn greater than1.0 indicates an increased probability for being found on the surface (Emini etal.,1985). IEDB was used to predict surface accessibility using default threshold value for each protein [35].

Epitopes antigenicity sites: Third steps of identification was antigenic sites for each protein with default threshold value was conducted using Kolaskar and Tongaonker antigenicity tool of IEDB [36]. This prediction is based on a semi-empirical approach, developed on physicochemical properties of amino acid residues and their frequencies of occurrence in experimentally known segmental epitopes and has the efficiency to detect antigenic peptides with about75%accuracy(threshold setting=1.000; KolaskarandTongaonkar,1990).

Hydrophobicity analysis: The amino acids making up the epitope are usually charged and hydrophilic in nature. Parker hydrophobicity scale was applied to evaluate the hydrophobicity. It is based on peptide retention time in high-performance liquid chromatography (HPLC) on reversed-phase column. Using a window of seven residues, these experimental values are calibrated for each of these residues and the arithmetical mean of the seven residue value was assigned to the fourth,(i +3), residue in the segment(threshold setting=1.678; Parkeretal.,1986).

### HLA binding peptide prediction

MHC epitope prediction: IEDB server (http://www.iedb.org) was used through specific tools to determine MHC1 and MHC II binding epitopes. This server uses specific scoring IC50 (inhibitory concentration 50) to predict epitopes that bind to different MHC class I and MHC class II alleles.

#### MHC class I epitope binding prediction

The MHC system is an example of receptor that can interact with linear ligands of variable lengths, and the method of prediction for MHC class I affinity was been tested on large set of quantitative peptide MHC class I measurement affinity on the IEDB, by using artificial neutral network (ANN) method and length of nine amino acid, all conserved epitopes bound with score equal or less than 300 IC50 for all three structural proteins were chosen for further analysis [37].

#### MHC class II epitope binding prediction

One of the major goals of the immunological research is the identification of MHC class II restricted peptide epitopes, and for that reason many computational tools have been developed but also their performance has lack of large scale systematic evaluation, and we use comprehensive dataset consisted from thousands of previously unpublished MHC peptide binding affinities and peptide MHC crystal structures, which all tested for CD4+ T cell responses to evaluate the publicly available MHC class II binding prediction tools performances. Therefore MHCII binding tool from IEDP was used by applying NN align as prediction method. Then using IC50 prediction value equal or less than 1000, all conserved epitopes were chosen for more analysis [38].

### Population Coverage

The estimation of the population coverage are based on the MHC binding with or without T cell restriction data, there for Nemours based tool was developed to predict population coverage of T cell epitope-based diagnostic and vaccines based on MHC binding with or without T cell restriction data. All alleles that interact with epitopes from 1,6 Fructose-bisphosphate aldolase sequences were subjected population coverage tool of IEDB (http://tools.iedb.org/tools/population/iedb_input) to calculate the whole world population coverage of MHC class I, MHC II and combined MHC I and II alleles for each protein [39].

### Homology Modelling

The 3D structure for the three different proteins was obtained using Raptor X structure prediction server [40] and Chimera 1.8 [41] was used to demonstrate the structure of proposed B cell and T cell epitopes that can be utilized for vaccine development.

## RESULTS

### Multiple Sequence Alignment

The conserved regions and amino acid composition for the reference sequence of *Schistosoma Mansoni* putative FBA are illustrated in figure 1 and 2 respectively. Alanine and Leucine were the most frequent amino acids (Table 2)

**Figure 1:**
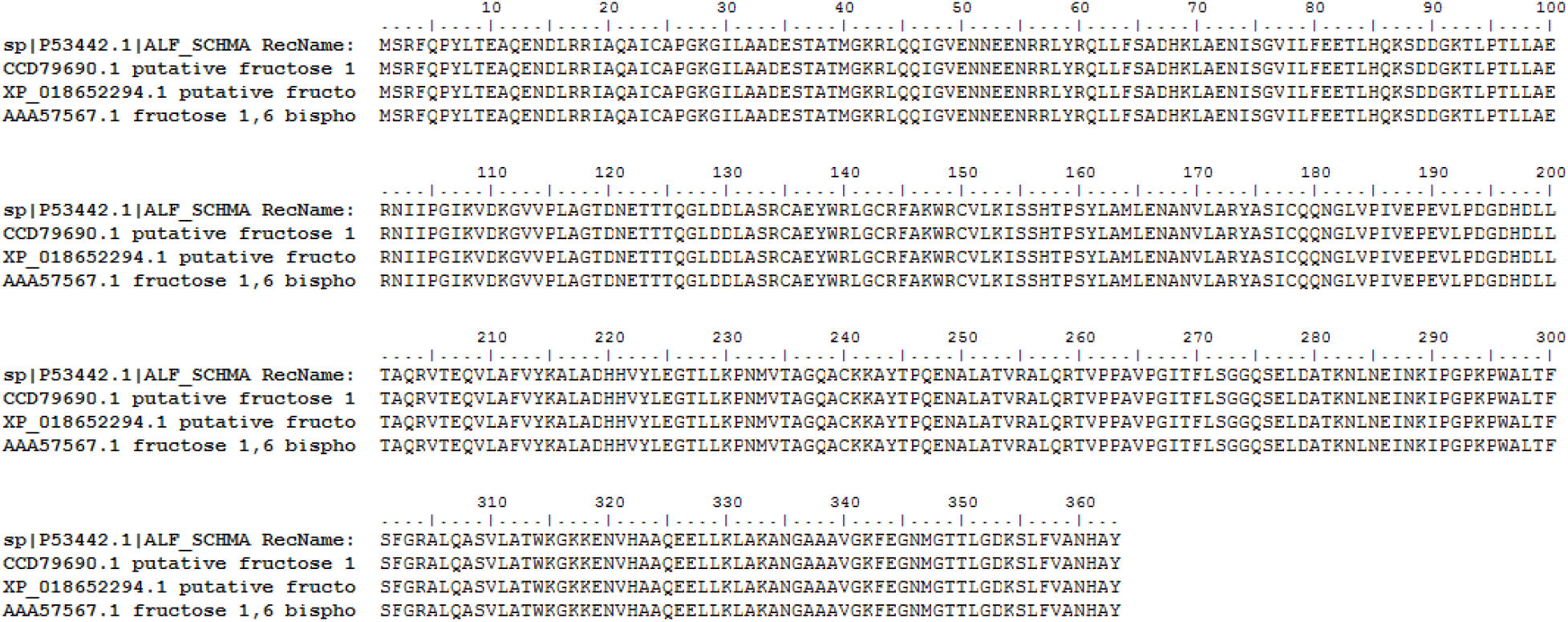
Multiple Sequence Alignment using BioEdit software.

**Figure 2:**
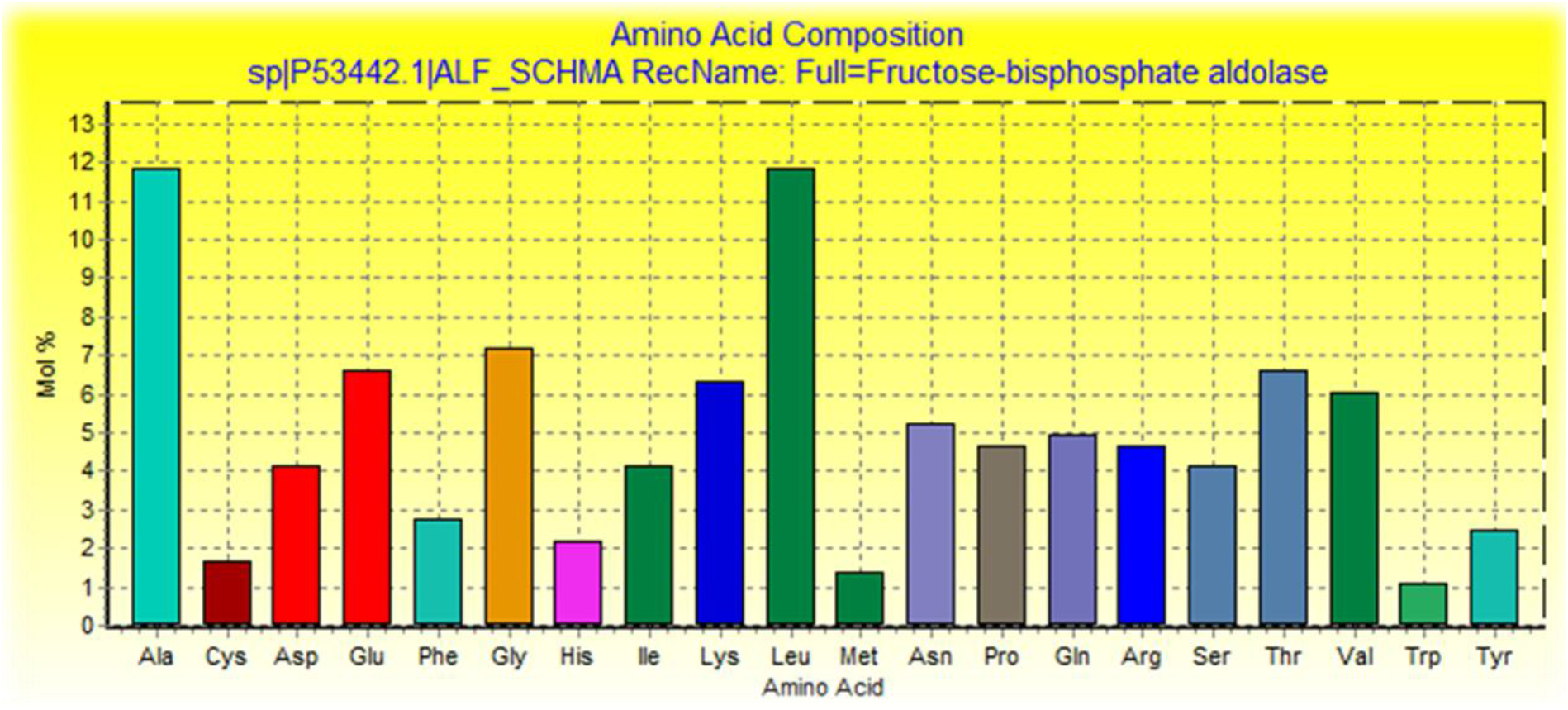
Amino acid composition for *Schistosoma Mansoni* FBA using BioEdit software.

**Table 2:** Molecular weight and amino acid frequency distribution of the protein

| Amino | Number | Mol% |
| --- | --- | --- |
| Ala | 43 | 11.85 |
| Cys | 6 | 1.65 |
| Asp | 15 | 4.13 |
| Glu | 24 | 6.61 |
| Phe | 10 | 2.75 |
| Gly | 26 | 7.16 |
| His | 8 | 2.2 |
| Ile | 15 | 4.13 |
| Lys | 23 | 6.34 |
| Leu | 43 | 11.85 |
| Met | 5 | 1.38 |
| Asn | 19 | 5.23 |
| Pro | 17 | 4.68 |
| Gln | 18 | 4.96 |
| Arg | 17 | 4.68 |
| Ser | 15 | 4.13 |
| Thr | 24 | 6.61 |
| Val | 22 | 6.06 |
| Trp | 4 | 1.1 |
| Tyr | 9 | 2.48 |
| Tyr | 9 | 2.48 |

### B-cell epitope prediction

The reference sequence of *Schistosoma Mansoni* FBA was subjected to Bepipred linear epitope 2, Emini surface accessibility, Kolaskar & Tongaonkar antigenicity and Parker hydrophilicity prediction methods to test for various immunogenicity parameters (Table 3 and Figures 3–6). Three epitopes have successfully passed the three tests. Tertiary structure of the proposed B cell epitopes was shown (Figure 7–9)

**Figure 3:**
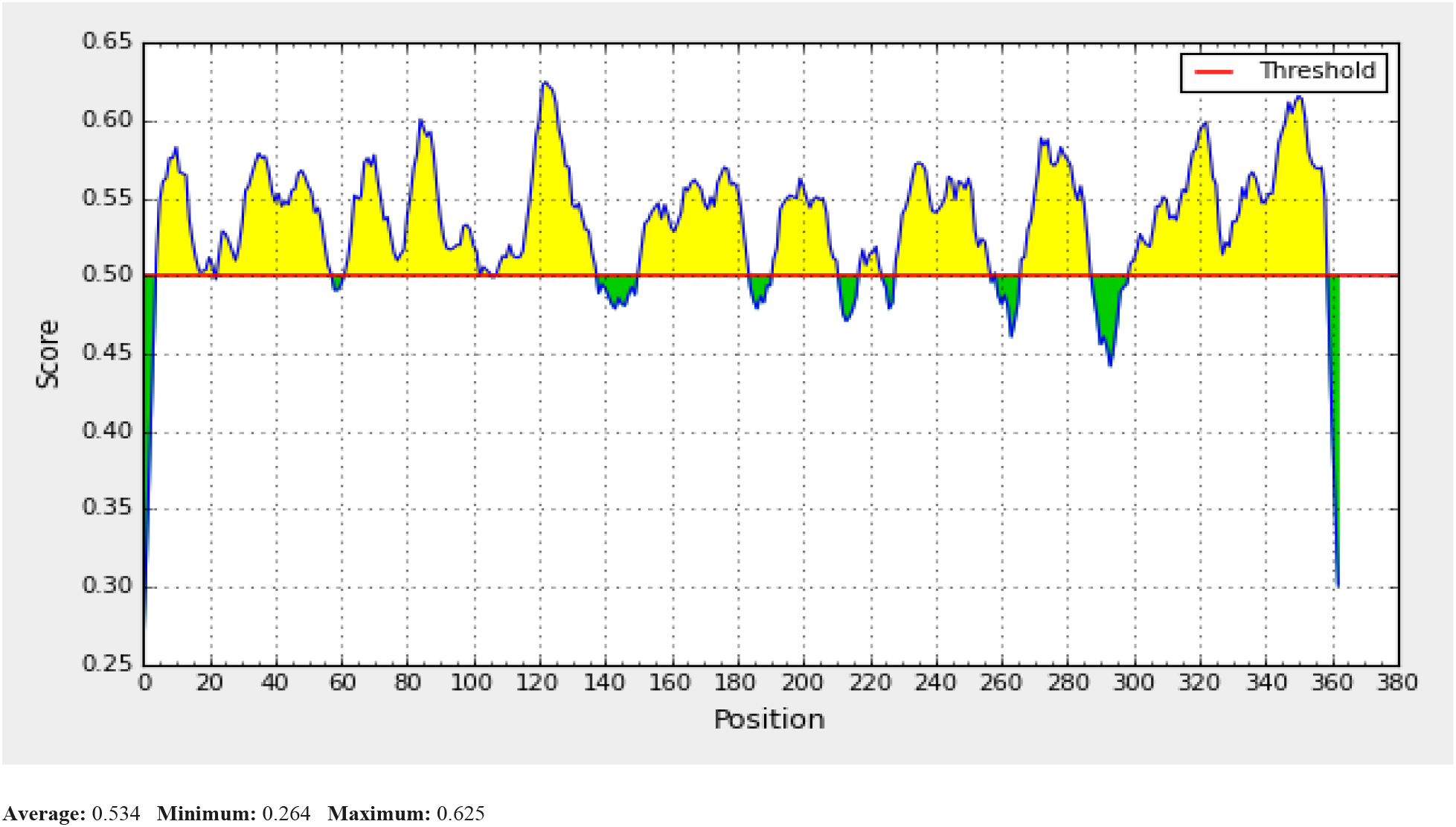
Bepipred Linear Epitope Prediction; Yellow areas above threshold (red line) are proposed to be a part of B cell epitopes and the green areas are not

**Figure 4:**
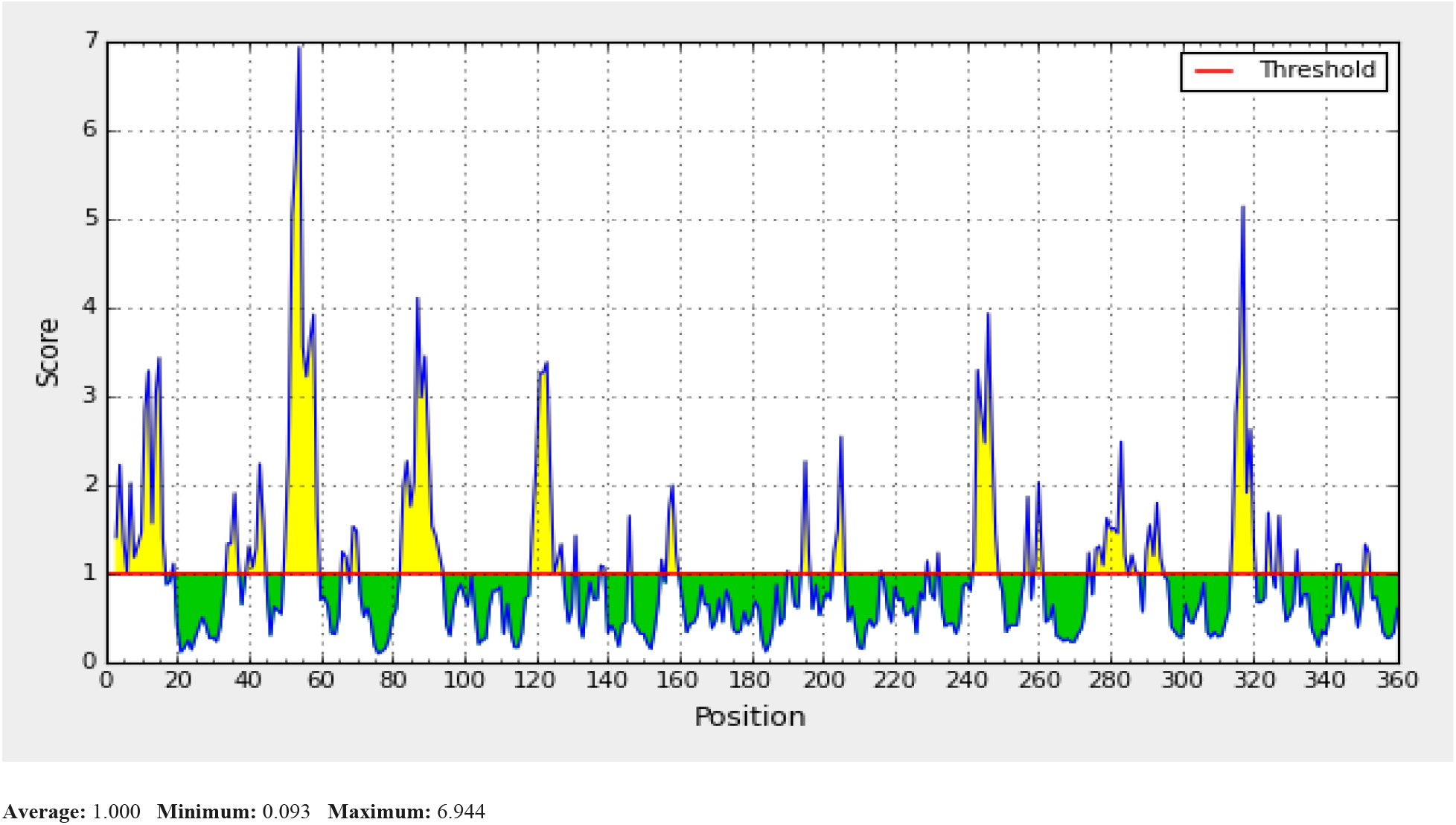
EMINI surface accessibility prediction; Yellow areas above the threshold (red line) are proposed to be a part of B cell epitopes and the green areas are not.

**Figure 5:**
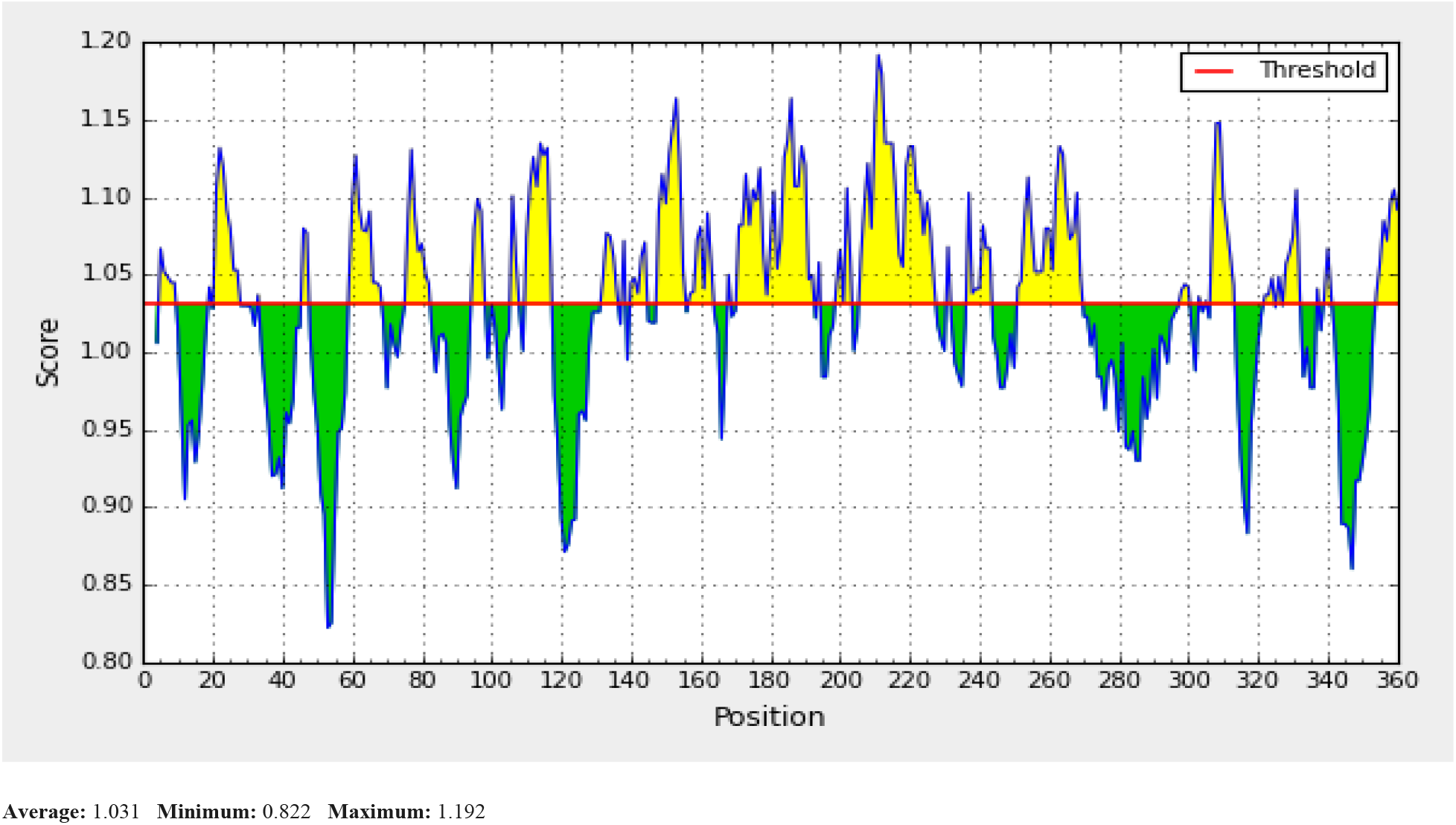
Kolaskar and Tonganokar antigenicity prediction; Yellow areas above the threshold (red line) are proposed to be a part of B cell epitopes and green areas are not.

**Figure 6:**
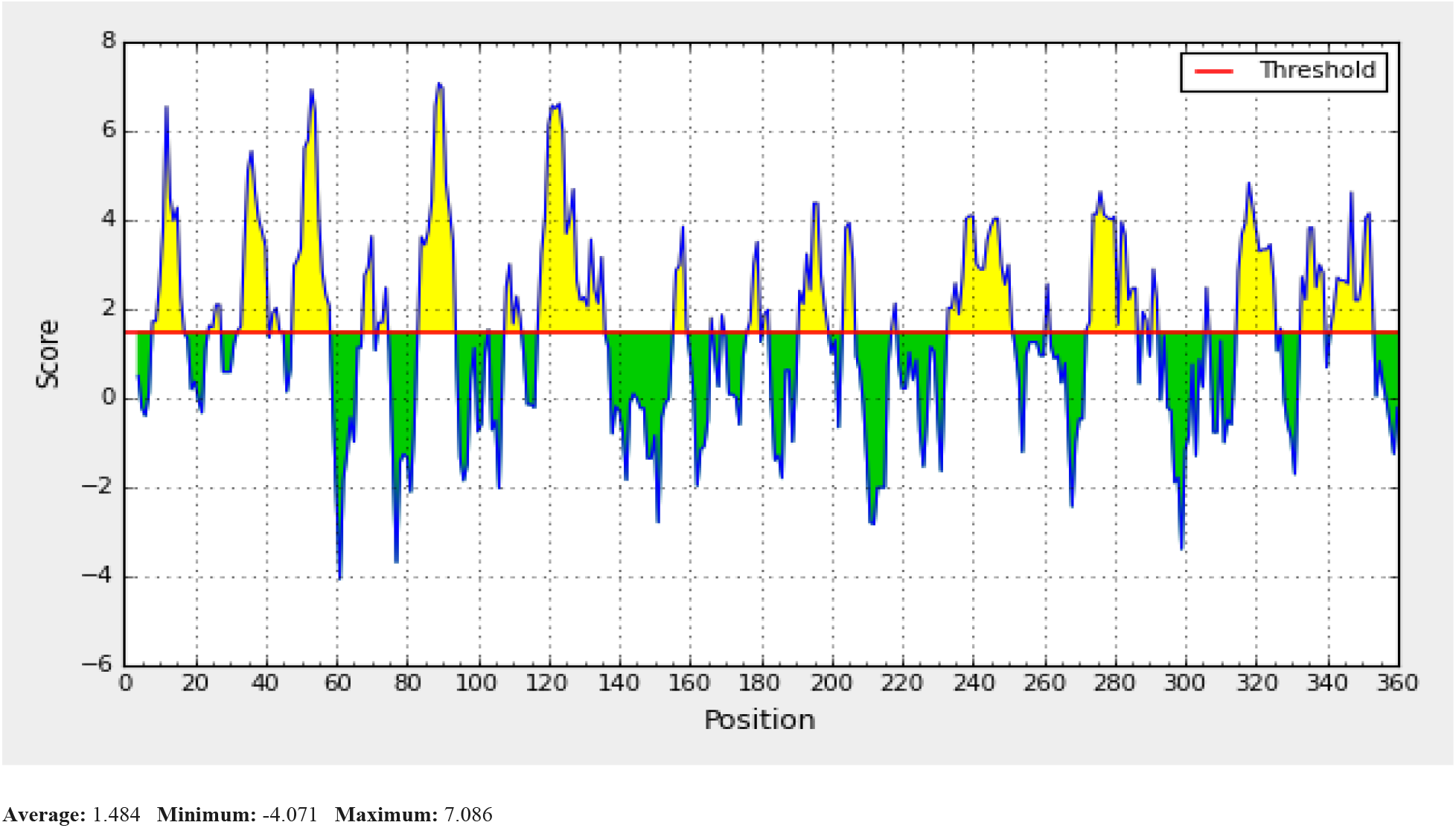
Parker Hydrophilicity prediction; Yellow areas above the threshold (red line) are proposed to be a part of B cell epitopes and green areas are not

**Figure 7:**
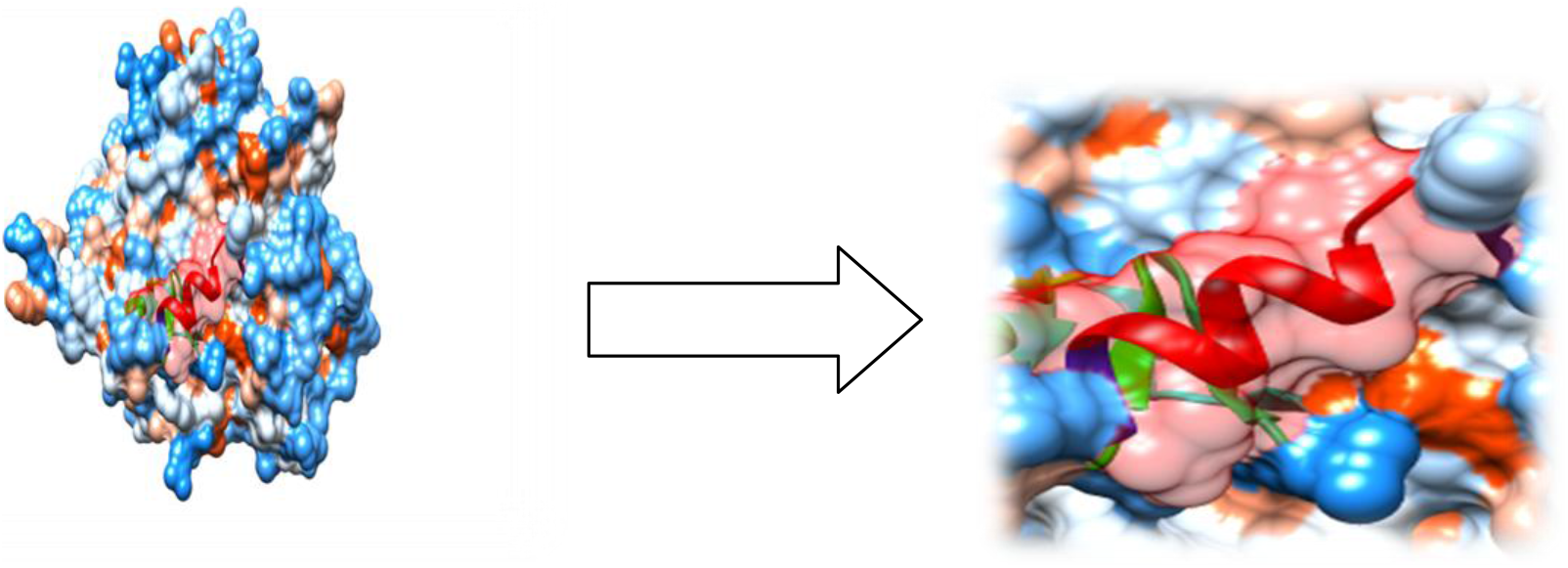
Proposed B cell epitope. The arrow shows position of **RRIAQAICA** with red colour at structural level using Chimera software.

**Figure 8:**
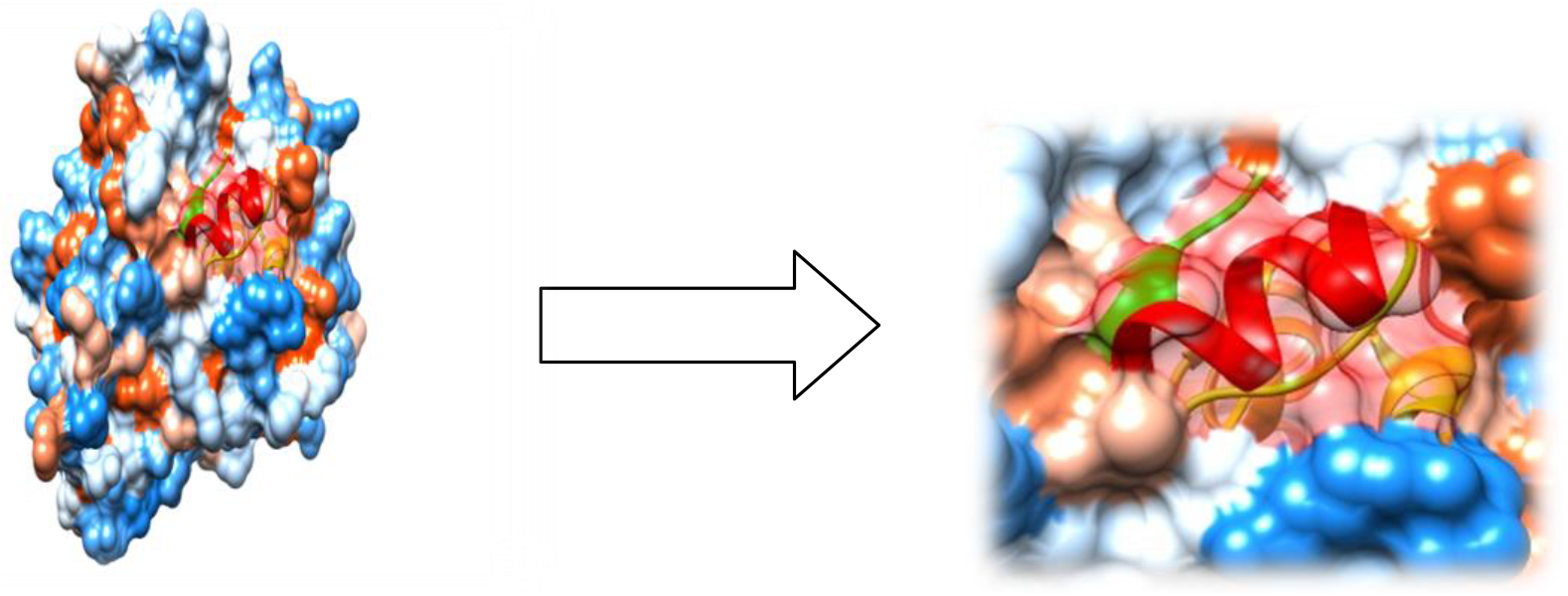
Proposed B cell epitope. The arrow shows position of **AQRVTEQV** with red colour at structural level using Chimera software.

**Figure 9:**
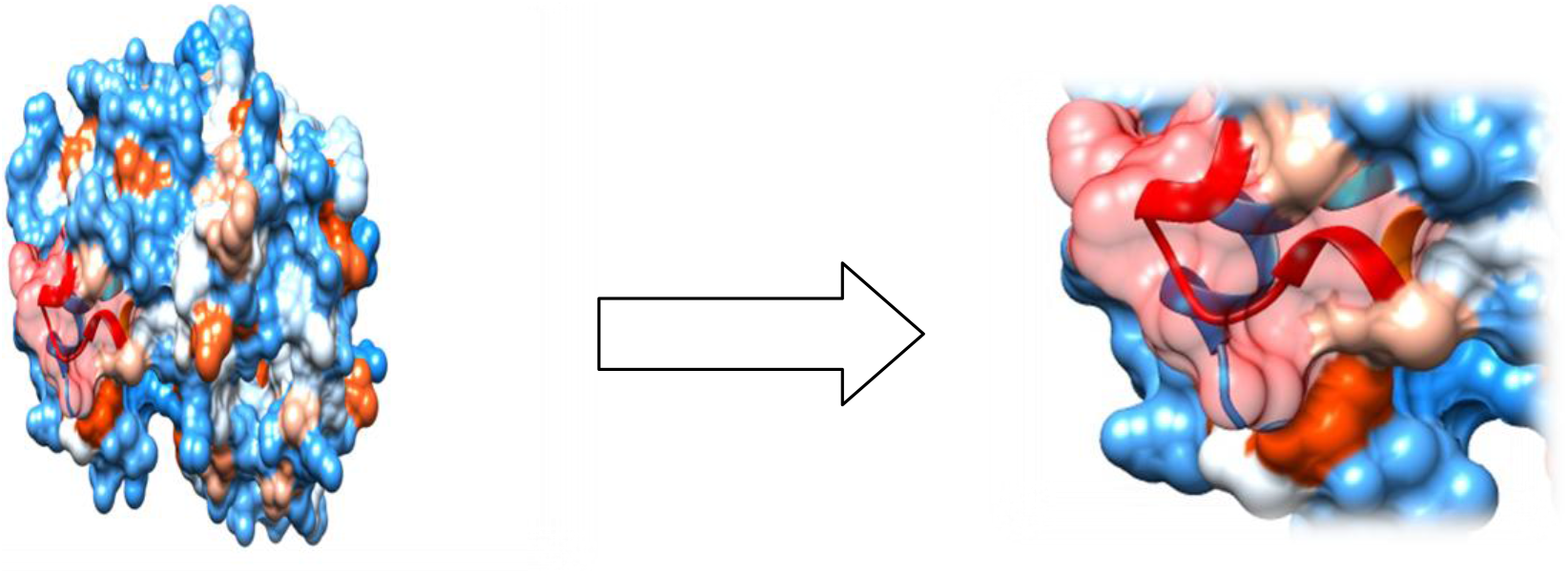
Proposed B cell epitope. The arrow shows position of **TWKGKKENV** with red colour at structural level using Chimera software.

**Table 3:**
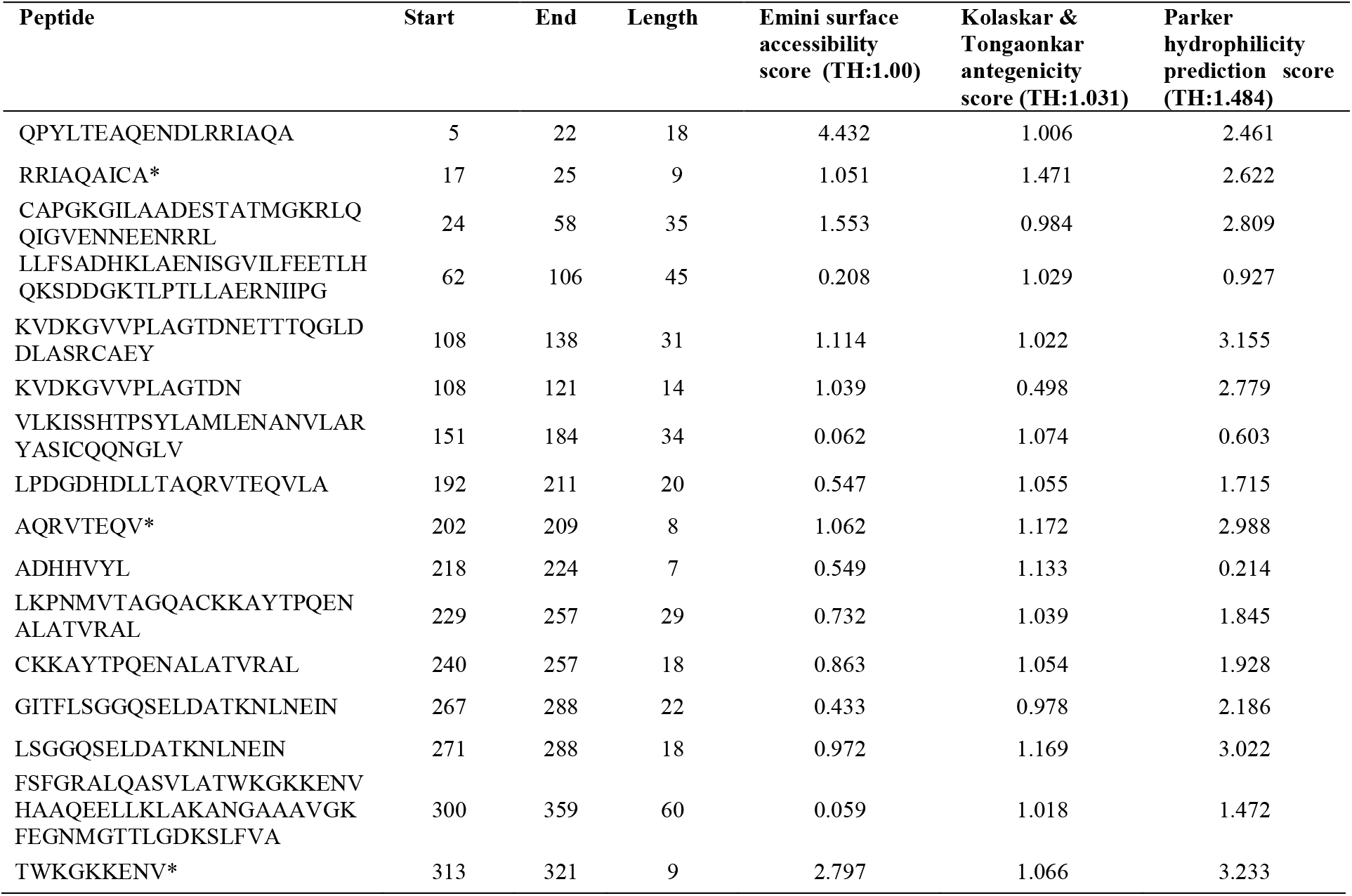
List of conserved peptides with their EMINI surface accessibility, Kolaskar & Tongaonkar antigenicity and Parker hydrophilicity scores. (*) Epitopes that successfully passed the three tests

### T-cell epitope prediction MHC I

The reference sequence was analyzed using (IEDB) MHC-1 binding prediction tool to predict T cell epitopes interacting with different types of MHC Class I alleles, based on Artificial Neural Network (ANN) with half maximal inhibitory concentration (IC50) <100 nm. 54 peptides were predicted to interact with different MHC1alleles. The top of five epitopes (most promising) and their corresponding MHC-1 alleles are shown in (Table 4) followed by the 3D structure of the proposed T cell epitope (Figure 10).

**Table 4:**
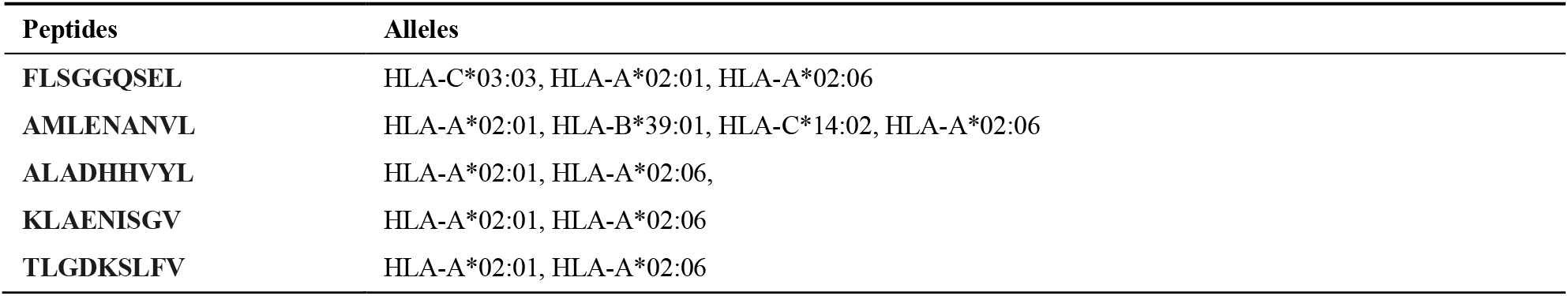
The top of five T cell epitopes and their corresponding MHC-1 alleles

**Figure 10:**
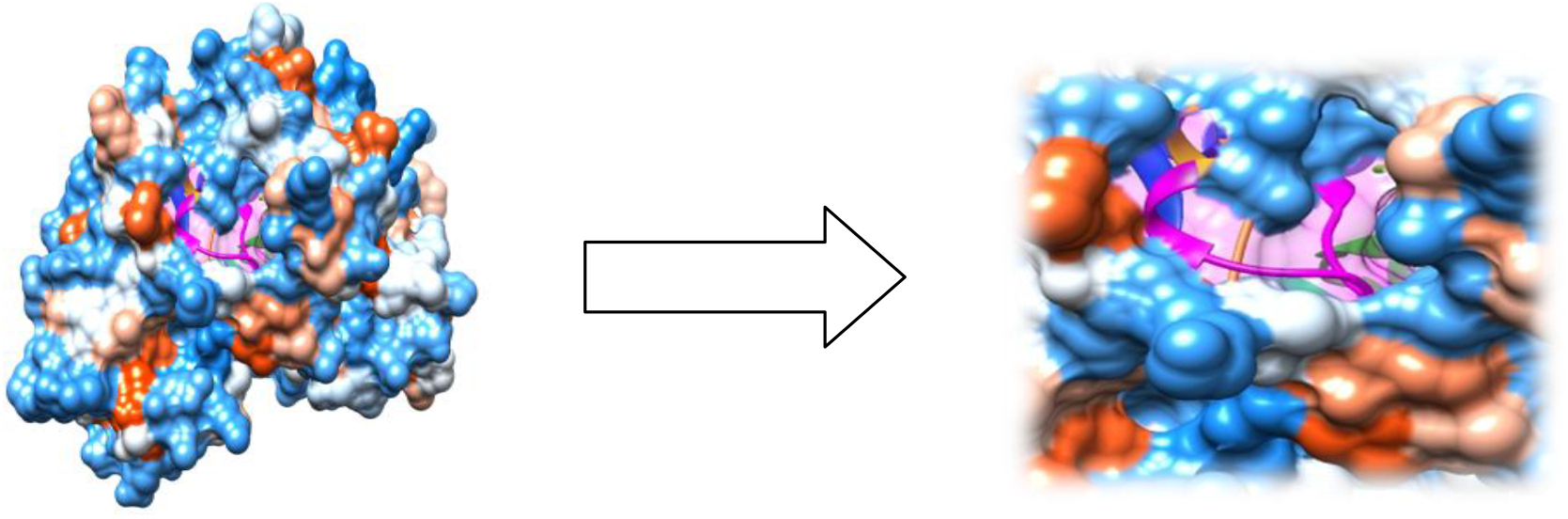
Proposed T cell epitopes that interact with MHC1. The arrow shows position of **FLSGGQSEL** with magenta colour at structural level using Chimera software.

### T-cell epitope prediction MHC II

Reference sequence was analyzed using (IEDB) MHC-II binding prediction tool based on NN-align with half maximal inhibitory concentration (IC50) <500 nm; there were 308 predicted epitopes found to interact with MHC-II alleles. The top five epitopes and their corresponding alleles are shown in (Table 4) along with the tertiary structure of the proposed epitope (Figure 10).

**Table 4:** The top of five T cell epitopes and their corresponding MHC-II alleles

| Peptide | MHC 2 alleles |
| --- | --- |
| LTFSFGRALQASVLA | HLA-DRB1*01:01, HLA-DRB1*09:01, HLA-DQA1*05:01/DQB1*03:01, HLA-DRB5*01:01, HLA-DRB1*04:05, HLA-DRB1*11:01, HLA-DRB1*07:01, HLA-DQA1*01:02/DQB1*06:02, HLA-DRB1*04:04, HLA-DRB1*04:01, HLA-DPA1*02:01/DPB1*01:01, HLA-DPA1*01:03/DPB1*02:01, HLA-DRB1*15:01, HLA-DPA1*03:01/DPB1*04:02, HLA-DRB1*13:02, HLA-DRB1*08:02, HLA-DRB4*01:01, HLA-DRB1*03:01, HLA-DPA1*01/DPB1*04:01 |
| ALTFSFGRALQASVL | HLA-DRB1*01:01, HLA-DRB1*09:01, HLA-DQA1*05:01/DQB1*03:01, HLA-DRB5*01:01, HLA-DRB1*11:01, HLA-DRB1*04:05, HLA-DRB1*07:01, HLA-DQA1*01:02/DQB1*06:02, HLA-DPA1*01:03/DPB1*02:01, HLA-DRB1*04:01, HLA-DPA1*02:01/DPB1*01:01, HLA-DRB1*04:04, HLA-DRB1*13:02, HLA-DPA1*03:01/DPB1*04:02, HLA-DRB1*15:01, HLA-DRB1*08:02, HLA-DRB1*03:01, HLA-DPA1*01/DPB1*04:01, HLA-DRB4*01:01 |
| LYRQLLFSADHKLAEL | HLA-DRB1*04:05, HLA-DRB1*07:01, HLA-DRB1*03:01, HLA-DRB1*01:01, HLA-DRB1*04:04, HLA-DRB1*11:01, HLA-DPA1*02:01/DPB1*01:01, HLA-DRB1*04:01, HLA-DRB5*01:01, HLA-DRB1*15:01, HLA-DRB4*01:01, HLA-DPA1*01:03/DPB1*02:01, HLA-DRB1*13:02, HLA-DRB1*09:01, HLA-DPA1*01/DPB1*04:01, HLA-DPA1*02:01/DPB1*05:01, HLA-DPA1*03:01/DPB1*04:02, HLA-DRB3*01:01 |
| ISGVILFEETLHQKS | HLA-DRB1*04:01, HLA-DPA1*02:01/DPB1*01:01, HLA-DQA1*05:01/DQB1*02:01, HLA-DPA1*03:01/DPB1*04:02, HLA-DPA1*01:03/DPB1*02:01, HLA-DRB1*04:04, HLA-DRB4*01:01, HLA-DRB5*01:01, HLA-DRB1*15:01, HLA-DRB1*01:01, HLA-DPA1*02:01/DPB1*05:01, HLA-DQA1*03:01/DQB1*03:02, HLA-DRB1*03:01, HLA-DRB1*11:01, HLA-DPA1*01/DPB1*04:01, HLA-DRB1*04:05, HLA-DQA1*04:01/DQB1*04:02, HLA-DRB1*07:01 |
| YRQLLFSADHKLAEN | HLA-DRB1*03:01, HLA-DRB1*04:05, HLA-DRB1*07:01, HLA-DRB1*01:01, HLA-DRB1*11:01, HLA-DRB1*04:04, HLA-DPA1*02:01/DPB1*01:01, HLA-DRB1*04:01, HLA-DRB5*01:01, HLA-DRB1*15:01, HLA-DRB4*01:01, HLA-DPA1*03:01/DPB1*04:02, HLA-DRB1*13:02, HLA-DRB1*09:01, HLA-DRB3*01:01, HLA-DPA1*02:01/DPB1*05:01, HLA-DPA1*01/DPB1*04:01 |

### Analysis of population coverage

All MHC I and MHC II epitopes were assessed for population coverage against the whole world using IEDB population coverage tool. For MHC 1, epitopes with highest population coverage were FLSGGQSEL (45.42%) and AMLENANVL (43.99%) as shown in (Figure 11 and Table 5). For MHC class II, the epitopes that showed highest population coverage were LTFSFGRALQASVLA (81.94%) and ALTFSFGRALQASVL (81.94%) (Table 6). When combined together, the epitopes that showed highest population coverage were LTFSFGRALQASVLA (81.94%), ALTFSFGRALQASVL (81.94%) and LYRQLLFSADHKLAE (80.93%) (Figure 12 and Table 7)

**Figure 11:**
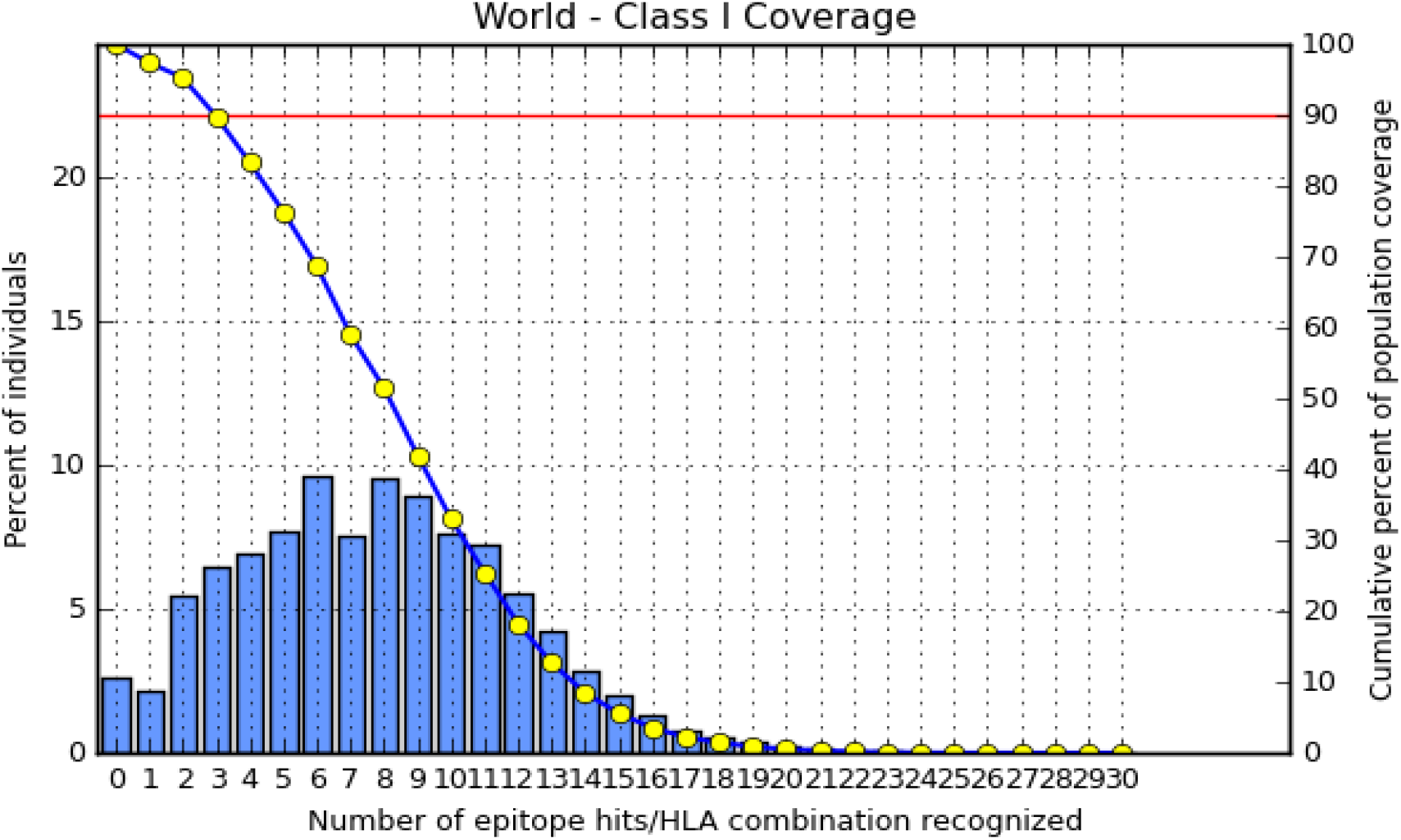
Population coverage for MHC class I epitopes.

**Figure 12:**
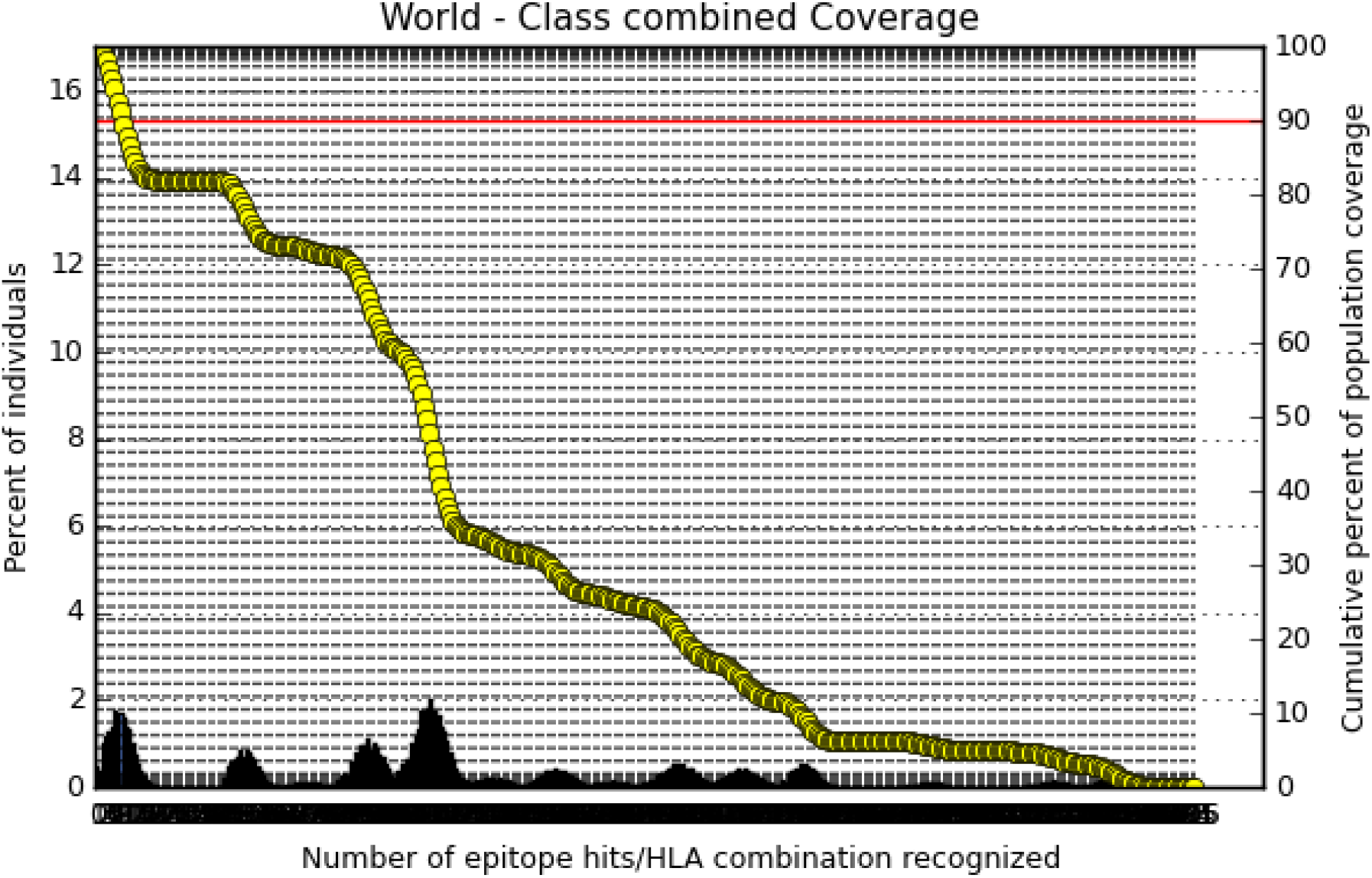
Population coverage for combined MHC class I & II epitopes.

**Table 5:** Population coverage of proposed peptides interaction with MHC class 1

| Epitope | Coverage Class I (%) | Total HLA hits |
| --- | --- | --- |
| FLSGGQSEL | 45.42% | 3 |
| AMLENANVL | 43.99% | 4 |
| ALADHHVYL | 40.60% | 2 |
| KLAENISGV | 40.60% | 2 |

**Table 6:** Population coverage of proposed peptides interaction with MHC class 2

| Epitope | Coverage Class II (%) | Total HLA hits |
| --- | --- | --- |
| LTFSFGRALQASVLA | 81.94% | 19 |
| ALTFSFGRALQASVL | 81.94% | 19 |
| LYRQLLFSADHKLAEL | 80.93% | 18 |
| YRQLLFSADHKLAEN | 80.93% | 17 |
| RQLLFSADHKLAENI | 79.58% | 15 |

**Table 7:** Population coverage of proposed peptide interaction with MHC class 1&2 combined

| Epitope | Coverage Class 1&2 combined (%) | Total HLA hits |
| --- | --- | --- |
| LTFSFGRALQASVLA | 81.94% | 19 |
| ALTFSFGRALQASVL | 81.94% | 19 |
| LYRQLLFSADHKLAEL | 80.93% | 18 |
| YRQLLFSADHKLAEN | 80.93% | 17 |
| RQLLFSADHKLAENI | 79.58% | 15 |
| TFSFGRALQASVLAT | 78.93% | 17 |
| VLKISSHTPSYLAML | 78.44% | 11 |
| ISGVILFEETLHQKS | 74.67% | 18 |
| RRLYRQLLFSADHKL | 73.11% | 17 |
| WALTFSFGRALQASV | 73.11% | 17 |

**Table 8:** The population coverage of whole world for the epitope set for MHC I, MHC II and MHC I&II combined.

| Country | MHC I | MHC II | MHC I&II combined |
| --- | --- | --- | --- |
| World | 97.4% | 48.63%* | 99. 53 %* |

In population coverage analysis of MHC II; thirteen alleles were not included in the calculation, therefore the above (*) percentages are for epitope sets excluding: HLA-DQA1*05:01/DQB1*03:01, HLA-DQA1*01:02/DQB1*06:02, HLA-DPA1*01:03/DPB1*02:01, HLA-DPA1*01/DPB1*04:01, HLA-DPA1 *02:01/DPB 1*05:01, HLA-DQA1*03:01/DQB1*03:02, HLA-DRB3*01:01, HLA-DRB4*01:01, HLA-DRB5*01:01, HLA-DQA1*04:01/DQB1*04:02, HLA-DPA1*03:01/DPB1*04:02, HLA-DQA1 *05:01/DQB 1*02:01, HLA-DPA1*02:01/DPB 1*01:01

## DISCUSSION

In this study, the author aimed to determine the highly potential immunogenic epitope by analysing putative fructose 1,6-bisphosphate aldolase for all *Schistosoma Mansoni* strains to develop a candidate vaccine. *Schistosoma Mansoni* putative fructose 1,6-bisphosphate aldolase reference sequence was subjected to Bepipred linear epitope prediction tests namely Emini surface accessibility test, Kolaskar and Tongaonkar antigenicity test and Parker hydrophilicity test, to determine the binding to B cell and to test the immunogenicity and hydrophilicity respectively. The epitopes should get above threshold scores for all prediction methods. Only eleven peptides were predicted by the IEDB to have probability of activating B cells. Only three epitopes succeeded Emini surface accessibility, Kolaskar and Tongaonkar antigenicity and Parker Hydrophilicity test, **RRIAQAICA** with scores 1.051, 1.471 and 2.622 respectively, **AQRVTEQV** with scores 1.062, 1.172 and 2.988 respectively, and **TWKGKKENV** with scores 2.797, 1.066 and 3.233. In spite of one linear epitope **QPYLTEAQENDLRRIAQA** have highest score with 4.432 for Emini surface accessibility, **RRIAQAICA** have highest score with 1.471 for Kolaskar and Tongaonkar antigenicity and **TWKGKKENV** have highest score with 3.233 for Parker Hydrophilicity prediction. This proves the remarkable stimulation, surface area and “water-loving” B cell epitopes.

By using IEDB, MHC I & II binding prediction tools the reference sequence was analysed to predict T cell epitopes. Fifty four conserved T cell epitopes were predicted to interact with MHC I alleles with IC50 < 100. Five of them were most promising and have high affinity to bind to the highest number of MHC I (FLSGGQSEL, AMLENANVL, ALADHHVYL, KLAENISGV and TLGDKSLFV). 308 predicted epitopes were interacted tightly with MHC II alleles with IC50 <500. Five of them were most promising and have high affinity to bind to the highest number of MHC II alleles (LTFSFGRALQASVLA, ALTFSFGRALQASVL, LYRQLLFSADHKLAE, YRQLLFSADHKLAEN, RQLLFSADHKLAENI). The best epitope with the highest population coverage for MHC I was FLSGGQSEL with 45.42% with three HLA hits, and coverage population for whole MHC I epitopes was 97.4%. The best epitopes with the highest population coverage for MHC II were LTFSFGRALQASVLA and ALTFSFGRALQASVL with 81.94% over nineteen HLA hits, and coverage population for whole MHC II epitopes was 48.63%.

One of the limitations of this study was the number of Fructose-bisphosphate aldolase (FBA) Sequences. Unfortunately, there were only four sequences retrieved from NCBI Database. Another limitation was the exclusion of certain HLA alleles from the MHC II and combined population coverage calculation.

Vaccination is the method of introducing antigenic agents to stimulate the immune system and to develop adaptive immunity to a specific disease. On other hand, immunoinformatics approaches focus mainly on the design and study of algorithms for mapping potential B- and T-cell epitopes, which speeds up the time and lowers the cost needed for laboratory analysis of pathogen gene products. [42]

## CONCLUSION

Vaccination is a method to protect and minimize the possibility of infection. Design of vaccines using insilico prediction methods is highly appreciated due to the significant reduction in cost, time and effort. Peptide vaccines overcome the side effects of conventional vaccines. We presented different peptides that can produce antibodies against FBA of *Schistosoma Mansoni* for the first time.

